# Exploring the consequences of redirecting an exocytic Rab onto endocytic vesicles

**DOI:** 10.1101/2023.02.09.527811

**Authors:** Xia Li, Dongmei Liu, Eric Griffis, Peter Novick

## Abstract

Bidirectional vesicular traffic links compartments along the exocytic and endocytic pathways. Rab GTPases have been implicated in specifying the direction of vesicular transport because anterograde vesicles are marked with a different Rab than retrograde vesicles. To explore this proposal, we sought to redirect an exocytic Rab, Sec4, onto endocytic vesicles by fusing the catalytic domain of the Sec4 GEF, Sec2, onto the CUE localization domain of Vps9, a GEF for the endocytic Rab, Ypt51. The Sec2GEF-GFP-CUE construct was found to localize to bright puncta predominantly near sites of polarized growth and this localization was strongly dependent upon the ability of the CUE domain to bind to the ubiquitin moieties added to the cytoplasmic tails of proteins destined for endocytic internalization. Sec4 and Sec4 effectors were recruited to these puncta with varying efficiency. The puncta appeared to consist of clusters of 80 nm vesicles and although the puncta are largely static, FRAP analysis suggests that traffic into and out of these clusters continues. Cells expressing Sec2GEF-GFP-CUE grew surprisingly well and secreted protein at near normal efficiency, implying that Golgi derived secretory vesicles were delivered to polarized sites of cell growth, where they tethered and fused with the plasma membrane despite the misdirection of Sec4 and its effectors. In total, the results suggest that while Rabs play a critical role in regulating vesicular transport, cells are remarkably tolerant of Rab misdirection.

---

Organelles along membrane transport pathways are linked by extensive, bidirectional, vesicular traffic. A key question is how the direction of any given vesicle is determined. Integral membrane proteins required for membrane fusion, such as the vesicle SNAREs, cycle continuously through both anterograde and retrograde vesicles (Lewis et al., 2000) and therefore appear to be ill suited to specify the direction of vesicle traffic (Whyte and Munro, 2002). In contrast, Rab GTPase activation and membrane association is reversible, with different Rabs tagging anterograde and retrograde vesicles. Thus Rabs appear better equipped to specify the direction of vesicle transport (Mizuno-Yamasaki et al., 2012). Rab activation requires a guanine nucleotide exchange factor (GEF) and Rab inactivation by GTP hydrolysis requires a GTPase activating protein (GAP). GDP-bound, lipid-anchored Rabs are subject to extraction from membranes by Rab GDP dissociation inhibitor (GDI) (Araki et al., 1990). By locating a GEF on one compartment and a GAP on another compartment, an asymmetric, unidirectional Rab distribution can be established. Rab effectors are defined by their recognition of a specific GTP-bound Rab and constitute a diverse collection of molecules that include molecular motors that drive the vectorial delivery of an organelle or vesicular carrier along a cytoskeletal track, tethers that mediate the initial recognition of the target compartment by a vesicular carrier and SNARE regulators that control the assembly of specific SNARE complexes and their function in membrane fusion (Grosshans et al., 2006b). As membrane flows along the exocytic or endocytic pathways, the Rabs associated with that membrane change, and as each new Rab decorates the membrane, it recruits a distinct set of effectors that help to define the functional identity of the membrane with which it is associated.

Rabs represent the largest branch of the GTPase superfamily, with more than sixty members in mammals and ten in yeast (Seabra et al., 2002). Reflecting the importance of Rabs as key nodes in the regulation of membrane traffic, a wide variety of human diseases have been attributed to defects in Rab expression, Rab prenylation, Rab GDI, Rab GEFs, Rab GAPs and Rab effectors (Hutagalung and Novick; Hutagalung and Novick, 2011; Seabra et al., 2002). Furthermore, a number of clinically important human pathogens exploit and disrupt our Rab regulatory pathways to promote their own intracellular agenda and to evade host defenses (Hutagalung and Novick, 2011).

Different Rabs are often networked to one another through their GEFs, GAPs and effectors, thereby generating regulatory circuits that can coordinate the various biochemical steps of each individual stage of membrane traffic and link together the stages of an entire transport pathway (Mizuno-Yamasaki et al., 2012). GEF and GAP cascades have been identified in which one Rab recruits the GEF that activates the downstream Rab and/or the GAP that inactivates the upstream Rab (Ortiz et al., 2002; Pusapati et al., 2012; Rana et al., 2015; Rivera-Molina and Novick, 2009; Suda et al., 2013; Zhu et al., 2009). These two cascade mechanisms can work in a counter-current fashion to direct a programmed series of Rab transitions as membrane flows along a membrane traffic pathway. In addition, positive feedback loops are formed through the ternary interactions of a Rab GEF, the activated Rab and one of its effectors (Horiuchi et al., 1997; Medkova et al., 2006). These positive feedback loops can function to sharpen Rab transitions and contribute to the maturation of carrier vesicles (Mizuno-Yamasaki et al., 2010; Stalder et al., 2013).

A key question concerns the importance of these regulatory mechanisms to membrane traffic in vivo. Prior efforts have disrupted individual regulatory elements (Nottingham et al., 2012; Ortiz et al., 2002; Pusapati et al., 2012; Rivera-Molina and Novick, 2009) or redirected a Rab regulatory component to an irrelevant target such as mitochondria (Blumer et al., 2013), but have not attempted to rewire a Rab regulatory circuit within the context of a functioning pathway and then assess the ramifications for membrane traffic. Here we seek to directly test the hypothesis that Rabs dictate the direction of vesicular transport. Our approach has been to fuse the catalytic GEF domain that activates the final Rab of the yeast exocytic pathway with a domain that normally serves to recruit the GEF for the first Rab of the endocytic pathway onto compartments of the endocytic pathway. Since the location of a GEF determines to a large extent where its substrate Rab can stably associate with membranes, this should have the effect of redirecting an exocytic Rab onto endocytic compartments.

Sec2, the exocytic GEF that we sought to redirect, activates Sec4, a Rab8 homolog that serves as the final Rab of the yeast secretory pathway (Walch-Solimena et al., 1997). The catalytic GEF domain near the amino terminus of Sec2 (aa’s 1-160) forms a homodimeric coiled-coil structure (Dong et al., 2007), while downstream regions are involved in its recruitment to Golgi-derived secretory vesicles through interactions with the upstream Rab Ypt32-GTP (Ortiz et al., 2002), a downstream effector Sec15 (Medkova et al., 2006) and the phosphoinositide PI(4)P (Mizuno-Yamasaki et al., 2010). Once activated, Sec4-GTP recruits Myo2, a type V myosin (Jin et al., 2011; Walch-Solimena et al., 1997). This leads to the movement of secretory vesicles along actin cables towards sites of polarized cell growth, including the tips of small buds and the necks separating large buds from the mother cell near the time of cytokinesis (Donovan and Bretscher, 2015). Sec4 also recruits the exocyst complex that tethers secretory vesicles to the plasma membrane (Guo et al., 1999; Salminen and Novick, 1989) and the SNARE regulator Sro7 that promotes exocytic fusion (Grosshans et al., 2006a).

To redirect the GEF domain of Sec2 we fused it to the CUE domain of Vps9, a GEF that activates Ypt51, a Rab5 homolog that serves as the first Rab of the endocytic pathway (Hama et al., 1999). The CUE domain of Vps9 binds to ubiquitin moieties that are added to the cytoplasmic tails of cell surface proteins destined for internalization into endocytic vesicles (Shideler et al., 2015). While the isolated CUE domain can act as a localization domain, recruitment of full length Vps9 might involve an additional interaction with Arf1-GTP (Nagano et al., 2019). Membrane recruitment of Vps9 is thought to occur at the Trans Golgi Network (TGN), which appears to play the role of the early endosome in yeast (Nagano et al., 2019) (Day et al., 2018). Vesicles carrying Vps9 and endocytic cargo bud from the TGN and are directed to the late endosome (Nagano et al., 2019). Activation of Ypt51 promotes delivery of endocytic cargo to late endosomes where cell surface proteins are deubiquitinated and incorporated into luminal vesicles through the action of the ESCRT complex, forming multi-vesicular bodies. Following fusion of the multi-vesicular bodies with the vacuole, the luminal vesicles are degraded. Here we examine the effects of expressing a fusion protein consisting of the GEF domain of Sec2 and the CUE domain of Vps9.

## Results

### Sec2GEF-GFP-CUE localizes to bright, focused puncta dependent upon the ubiquitin-binding activity of the CUE domain

To explore the effects of replacing the normal localization domain of Sec2 with that of Vps9, we fused the GEF domain (aa 1-160) of Sec2 to GFP and then to the CUE domain of Vps9 (aa 408-450) generating Sec2GEF-GFP-CUE. Unless otherwise indicated, we expressed this construct from the *ADH1* promoter as the sole copy of Sec2, with or without the M419D CUE domain mutation that blocks binding to ubiquitin (Shideler et al., 2015). As controls, we expressed full length Sec2 fused to GFP (Sec2-GFP) and the GEF domain alone fused to GFP (Sec2GEF-GFP). As previously shown (Elkind et al., 2000), Sec2-GFP exhibited concentrations at the tips of small buds or across the necks of large budded cells, reflecting its association with secretory vesicles (Fig. 1A). Sec2GEF-GFP, lacking the Ypt32, Sec15 and PI(4)P binding sites that serve to recruit Sec2 to secretory vesicles, was predominantly cytosolic, nonetheless about 30% of the cells exhibited detectable, bud tip or neck localization. This residual localization likely reflects its association with Sec4 on secretory vesicles as the percentage increased to about 50% upon Sec4 overexpression (Fig. 1B). Sec2GEF-GFP-CUE exhibited several bright, tightly focused puncta per cell as well as numerous dimmer puncta and little cytosolic background. The bright puncta were frequently near sites of polarized cell surface expansion. However, the puncta within buds were often closer to the sides of the buds rather than at the tips, and localization at necks was in the form of a focused punctum rather than a bar across the neck. Sec2GEF-GFP-CUE^M419D^ showed largely cytosolic localization with no concentration other than a slightly higher fluorescence of the nucleus relative to the cytoplasm (Fig. 1A). A minor fraction of cells expressing Sec2GEF-GFP-CUE^M419D^ exhibited polarized fluorescence and, as in the case of Sec2GEF-GFP, this percentage increased in cells overexpressing Sec4 (Fig. 1B). Together, these data indicate that the CUE domain serves to efficiently localize Sec2GEF-GFP-CUE and that this function relies on its ability to interact with ubiquitin. The localization is distinct from that of full length Sec2, nonetheless many structures bearing Sec2GEF-GFP-CUE localize relatively close to the normal sites of cell surface growth despite their abnormally punctate appearance. Similar results were observed with constructs employing mCherry or NeonGreen as fluorescent tags (see below).

**Figure 1.**
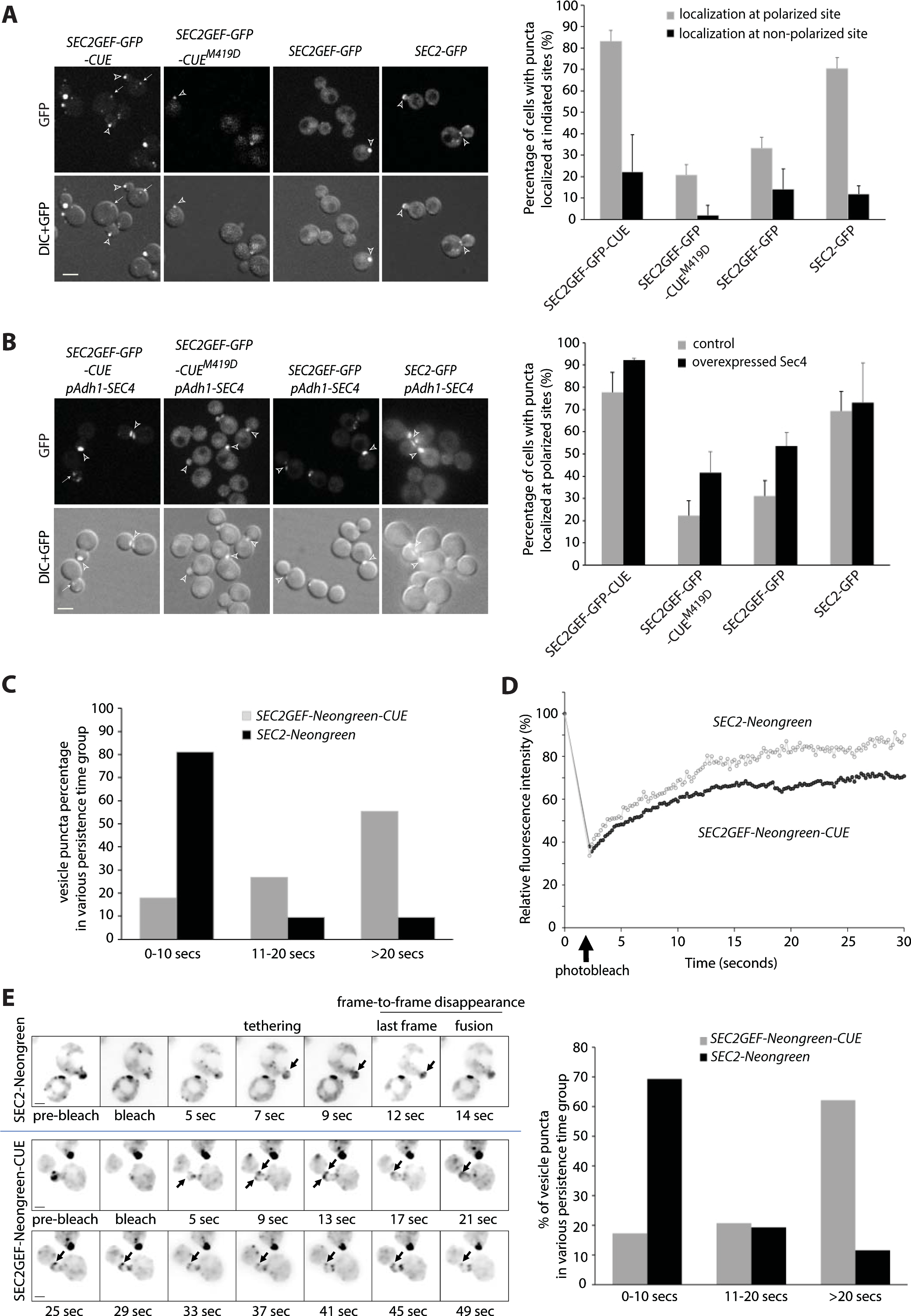
Sec2GEF-GFP-CUE and Sec2GEF-Neongreen-CUE localize to polarized sites with reduced dynamics and overexpression of Sec4 enhances the localization of Sec2GEF-GFP or Sec2GEF-GFP-CUE^M419D^ at polarized sites. **(A)** Shown are GFP images and DIC images overlaid with GFP images of representative cells grown to early log phase in SC medium at 25°C. The strains examined are *sec2Δ* containing an integrated plasmid expressing full length Sec2-GFP, Sec2GEF-GFP, Sec2GEF-GFP-CUE or Sec2GEF-GFP-CUE^M419D^ from the Adh1 promoter. Open arrowheads and arrows point to puncta localized at polarized sites including bud tip or bud neck and at non-polarized sites, respectively. Bars, 5μm. Sec2-GFP, Sec2GEF-GFP, Sec2GEF-GFP-CUE or Sec2-GEF-GFP-CUE^M419D^ puncta localization at polarized or non-polarized sites was quantified and shown on the right panel. The error bars in the graph are the SD from three independent experiments. **(B).** Representative images of GFP-label, mCherry-Sec4, merged and overlay DIC with merged images in *SEC2GEF-GFP-CUE, SEC2GEF-GFP-CUE^M419D^, SEC2GEF-GFP* or *SEC2-GFP* strains as indicated on the left panel. Bar, 5 μm. Open arrowheads and arrows point to GFP puncta localized at polarized sites and non-polarized sites, respectively. The localization of Sec2GEF-GFP-CUE, Sec2GEF-GFP-CUE^M419D^, Sec2GEF-GFP and control Sec2-GFP at polarized sites in absence or presence of overexpressed Sec4 was quantified and shown on the right panel. Sec2-GFP is well localized at polarized sites and Sec4 overexpression doesn’t affect it. **(C).** Early-log phase cells expressing Sec2-Neongreen or Sec2GEF-Neongreen-CUE as the sole copy of Sec2 under Adh1 promoter were analyzed by time-lapse fluorescence imaging for 90 seconds (see Movie S1 and Movie S2). The persistence time (in seconds) of identified vesicle puncta in wild-type *SEC2-Neongreen* cells (n=26) and mutant *SEC2GEF-Neongreen-CUE* cells (n=29) was determined as described in Materials and methods. Since the acquisition time is a total of 90 seconds and more than 50% of the puncta tracked in *SEC2GEF-Neongreen-CUE* cells persisted until the end of acquisition, we describe the vesicle dynamics using the percentage of vesicle puncta in various persistence time groups. **(D).** FRAP experiments showing the normalized fluorescence intensity in the entire small bud for tagged full length Sec2-Neongreen or Sec2GEF-Neongreen-CUE (see Movie S3 and Movie S4). The original fluorescence intensity (time = 0 sec) is normalized to 100% before photobleaching and during the time-lapse acquisition the cells selected are photobleached (time = 2sec, marked by the arrow). Plotted points represent the mean value of five replicate samples each. The photobleach and FRAP experimental details are described in *Materials and methods*. **(E).** Still frames of a typical vesicle tracking experiment after FRAP in wild-type *SEC2-Neongreen* or *SEC2GEF-Neongreen-CUE* cells. An inverted monochrome maximum projection is shown for clarity. Arrows point to Sec2-Neongreen or Sec2GEF-Neongreen-CUE puncta tethered to the membrane (See also Movie S5 and Movie S6). Bar, 2um. Using Kymograh to measure the persistence time (likely from tethering to fusion) of the vesicle puncta shown by the arrows in left panel. The quantification was done in wild-type *SEC2-Neongreen* cells (n=42) and mutant *SEC2GEF-Neongreen-CUE* cells (n=56), and the percentage of vesicle puncta in various persistence time groups are shown in right panel.

Movies were made using constructs containing the more photo stable NeonGreen variant. These revealed that the bright Sec2GEF-NeonGreen-CUE puncta were largely static, showing little movement over a 30 sec time frame. Dimmer puncta were more mobile but did not exhibit directed movement, and most persisted over the 30 sec observation period (Fig. 1C and supplemental movie S1). In contrast, Sec2-NeonGreen structures were more dynamic, and many smaller puncta displayed rapid, directed movement towards bud tips or bud necks. These persisted for an average of only 5 sec (Fig. 1C and supplemental movie S2). The dynamic localization of Sec2 reflects its association with secretory vesicles that fuse with the plasma membrane soon after delivery by Myo2 on actin cables (Donovan and Bretscher, 2015; Elkind et al., 2000).

While individual vesicles carrying fluorescently tagged proteins could be tracked, they became challenging to follow once they enter the bud due to the existing pool of vesicles at sites of polarized growth. Photobleaching all signals from existing vesicles in the bud allowed better visualization of new vesicles entering the bud from the mother cell. The experimental condition used for photobleaching the entire bud with a 405-nm laser did not affect cell growth, suggesting that the bleach event was not disturbing essential processes in exocytosis. After photobleaching the entire small bud, the rate of recovery of Sec2GEF-Neongreen-CUE was similar to that of Sec2-Neongreen, but less total signal was recovered in Sec2GEF-Neongreen-CUE cells (Fig. 1D and Supplemental movie S3, S4). Thus, although the bright Sec2GEF-GFP-CUE puncta appear static, additional Sec2GEF-GFP-CUE protein can be recruited to pre-existing puncta.

To quantitate the time of residence of individual vesicles at the plasma membrane prior to fusion we photobleached the bud and then followed delivery of vesicles from the unbleached region of the cell to the bud tip. We documented events in which the initial vesicle immobilization at the cell cortex was observed and the subsequent disappearance was seen in one of the three middle imaging planes thereby excluding events in which the vesicle moved out of the focal plane (Fig. 1E, left panel). Most vesicles carrying Sec2-NeonGreen (70%) remained stationary at the cell cortex for less than 10 sec before they disappeared (Fig. 1E and supplemental movie S5). In cells expressing Sec2GEF-Neongreen-CUE there were fewer puncta and 60% of them remained stationary for longer than 20 sec, suggesting a delay after tethering and prior to fusion (Fig. 1E and supplemental movie S6).

### Sec2GEF-CUE is efficiently recruited to the enlarged late endosomes in *vps4Δ* cells

Since the punctate localization of Sec2GEF-GFP-CUE was largely dependent upon the ubiquitin binding activity of its Vps9-derived CUE domain, we anticipated that it would co-localize with endogenous Vps9. About half of the Sec2GEF-GFP-CUE puncta were positive for Vps9-mCherry, regardless of their position within the cell, while Sec2-GFP displayed almost no co-localization with Vps9-mCherry (Supplemental Figure S1). The ubiquitin moiety on endocytic cargo usually remains accessible to the cytoplasm for only a short time before the cargo is deubiquitinated and incorporated into luminal vesicles in late endosomes by the ESCRT machinery (Shideler et al., 2015). The transient nature of CUE binding sites results in a high cytoplasmic background of Vps9-mCherry, which could explain the incomplete co-localization of Sec2GEF-GFP-CUE and Vps9-mCherry. We re-examined the co-localization in a *vps4Δ* mutant. In this ESCRT-defective mutant ubiquitinated cargo remains exposed to the cytoplasm on the surface of an enlarged late endosome, termed a “class E” compartment, leading to enhanced recruitment of Vps9 and a reduction in the cytoplasmic pool (Shideler et al., 2015). Sec2GEF-GFP-CUE localized to additional puncta at non-polarized sites in *vps4Δ* cells (Supplemental Figure S2A). Most of the Sec2GEF-GFP-CUE puncta at both polarized and non-polarized sites were also labeled with Vps9-mCherry, while Sec2-GFP structures were not positive for Vps9-mCherry (Fig. 2A). Vps9 serves to recruit and activate the Rab5 homolog, Ypt51 on endosomes (Hama et al., 1999). We observed extensive co-localization of mCherry-Ypt51 with Sec2GEF-GFP-CUE at both polarized and non-polarized sites in *vps4Δ* cells (Fig. 2B). No co-localization was observed with Sec2-GFP. We also observed substantial co-localization of the late endosome marker Vps8-mCherry with Sec2GEF-GFP-CUE, but not with Sec2-GFP in *vps4Δ* cells (Fig. 2C). Thus, the CUE domain serves to efficiently recruit Sec2GEF-GFP-CUE to endocytic membranes.

**Figure 2.**
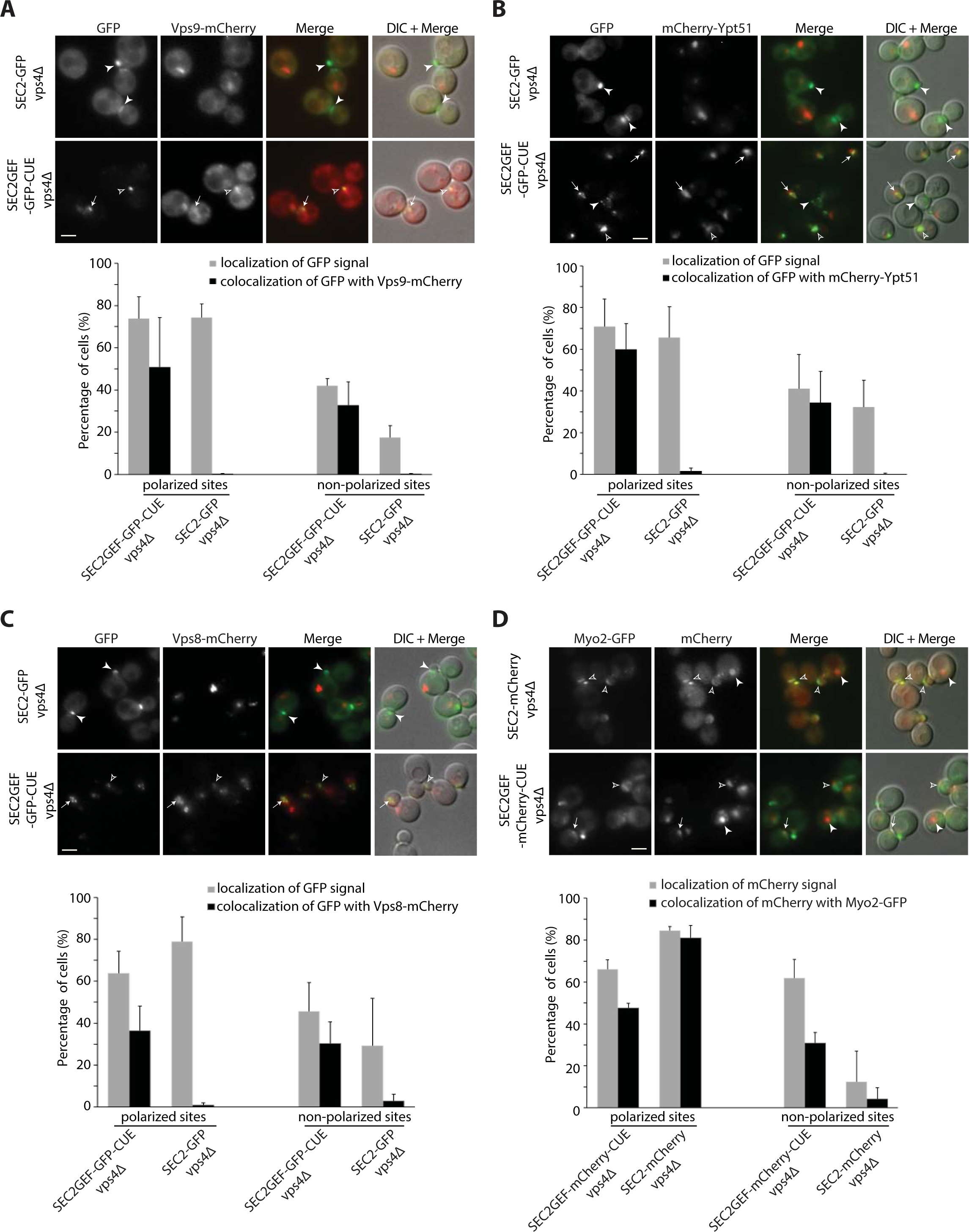
Localization of Sec2GEF-GFP-CUE (Sec2GEF-mCherry-CUE) or Sec2-GFP (Sec2-mCherry) and their colocalization with distinct Rabs or Rab effectors on the secretory and endocytic pathways in *vps4Δ* strains. **(A–C).** Live-cell fluorescence microscopy of *SEC2-GFP* and *SEC2GEF-GFP-CUE* strains expressing Vps9-mCherry (A), mCherry-Ypt51 (B) or Vps8-mCherry (C) are shown on upper panels. The quantification of Sec2-GFP and Sec2GEF-GFP-CUE localization at polarized or non-polarized sites and their colocalization with indicated mCherry-tagged proteins are shown in the lower panels. Open arrowheads and arrows indicate respectively polarized sites and non-polarized sites at which GFP colocalizes with mCherry-tagged proteins, while closed arrowheads point to GFP puncta which are not colocalized with the other protein examined. In all quantitation graphs the error bars represent the SD from three independent experiments. Scale bar, 5 μm. **(D).** Localization of Sec2GEF-mCherry-CUE or Sec2-mCherry and their colocalization with Myo2-GFP in *vps4Δ* strains. Upper panel shows representative GFP fluorescence, mCherry fluorescence, merged, and DIC overlaid with merged images. The quantification of Sec2-mCherry and Sec2GEF-mCherry-CUE localization at polarized or non-polarized sites and their colocalization with Myo2-GFP proteins are shown in lower panel. Open arrowheads and arrows respectively indicate polarized sites and non-polarized sites at which mCherry colocalizes with Myo2-GFP, while closed arrowheads point to mCherry puncta which are not colocalized with Myo2-GFP. In all quantitation graphs the error bars represent the SD from three independent experiments. Scale bar, 5 μm.

We also asked if the enlarged late endosomes bearing Sec2GEF-GFP-CUE or Sec2GEF-mCherry-CUE in *vps4Δ* cells recruit Sec4 and Sec4 effectors. We observed nearly complete co-localization of GFP-Sec4 and Sec2GEF-mCherry-CUE at both polarized or non-polarized sites in *vps4Δ* cells, while co-localization of GFP-Sec4 and Sec2-mCherry was mainly at polarized sites (Supplemental Figure S2B). There was extensive co-localization of Myo2-GFP and Sec2GEF-mCherry-CUE or Sec8-mCherry and Sec2GEF-GFP-CUE at polarized sites in *vps4Δ* cells (Fig. 2D and Supplemental Figure S2C). However, at non-polarized sites there was only partial co-localization of Myo2-GFP and Sec2GEF-mCherry-CUE (Fig. 2D) and no co-localization of Sec8-mCherry with Sec2GEF-GFP-CUE (Supplemental Figure S2C). No co-localization of the Golgi marker, Sec7-dsRed, was seen with either Sec2GEF-GFP-CUE or Sec2-GFP in *vps4Δ* cells (Supplemental Figure S2D). The observation that most of the compartments formed in response to Sec2GEF-CUE expression in *vps4Δ* cells bear both endosomal markers and secretory vesicle markers indicate that Sec2GEF-CUE is efficiently recruited to endosomal membranes in these cells and suggests that these compartments have a mixed identity.

Since the vesicle clusters marked by Sec2GEF-GFP-CUE had some characteristics of both an exocytic compartment and an endocytic compartment, we explored whether they would be marked by PI(4)P, the phosphoinositide made in the TGN, or PI(3)P, the phosphoinositide made in early endosomes. Neither Sec2-GFP nor Sec2GEF-GFP-CUE colocalized to a large degree with mCherry-FAPP1-PH, a probe for PI(4)P, in either a *VPS4* background or a *vps4Δ* background (Supplemental Fig S3 A and B). Sec2-GFP exhibited no colocalization with mRFP-FYVE(EEA1), a probe for PI(3)P in either a *VPS4* background or a *vps4Δ* background (Supplemental Fig S3 C and D). While Sec2GEF-GFP-CUE did not colocalize significantly with mRFP-FYVE(EEA1) in a *VPS4* background, about 30% of the Sec2GEF-GFP-CUE puncta did colocalize with mRFP-FYVE(EEA1) in a *vps4Δ* background. Colocalization was observed at both polarized and non-polarized sites (Supplemental Fig S3 C and D).

### Sec2GEF-CUE can recruit Sec4 and Sec4 effectors to polarized sites but is less efficient in the recruitment of effectors to non-polarized sites

Since we observed extensive co-localization of GFP-Sec4 and Sec2GEF-mCherry-CUE at both polarized or non-polarized sites in *vps4Δ* cells, we examined GFP-Sec4 in *VPS4* cells expressing Sec2GEF-mCherry-CUE or Sec2-mCherry. Nearly all Sec2GEF-mCherry-CUE puncta were also labeled with GFP-Sec4 whether they were near sites of polarized growth or at non-polarized sites. In control cells expressing Sec2-mCherry both proteins were found predominantly at polarized sites (Fig. 3A). These results are consistent with the observation that Rab GEFs can recruit their substrate Rabs to ectopic sites (Blumer et al., 2013).

**Figure 3.**
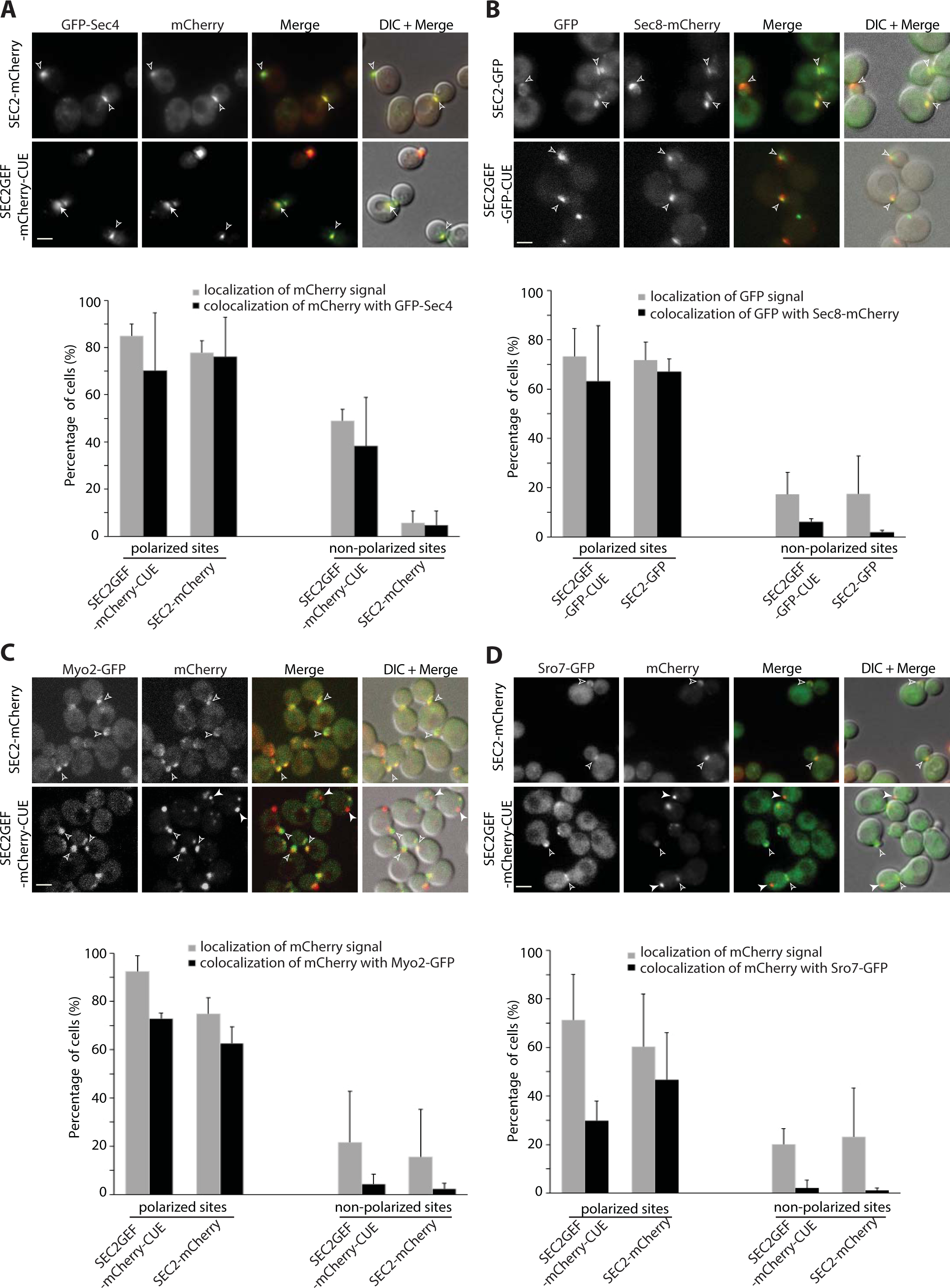
Sec2GEF-GFP-CUE recruits Sec4 to both polarized and non-polarized sites, but preferentially recruits Sec4 effectors to polarized sites. From Panel A to Panel D, representative images of the mCherry-fluorescence channel, GFP channel, merged image or overlay DIC with merged images of indicated cells are shown in the upper panels and quantification of the results are shown in the lower panels. The error bars represent the SD from three independent experiments. Scale bars, 5μm. (**A).** Shown are colocalization results of GFP-Sec4 with Sec2-mCherry or Sec2GEF-mCherry-CUE. Open arrowheads point to colocalized GFP-Sec4 with Sec2-mCherry or Sec2GEF-mCherry-CUE at polarized sites, bud tip in small buds or bud neck in large buds, respectively. Arrows point to colocalized GFP-Sec4 with Sec2GEF-mCherry-CUE at non-polarized sites. There is rarely colocalization of GFP-Sec4 with Sec2-mCherry at non-polarized sites. (**B).** Shown are colocalization results of exocyst Sec8-mCherry with Sec2-GFP or Sec2GEF-GFP-CUE. Open arrowheads point to colocalized Sec8-mCherry with Sec2-GFP or Sec2GEF-GFP-CUE at polarized sites. At non-polarized sites there is low percentage colocalization of Sec8-mCherry with Sec2GEF-GFP-CUE, and barely any with Sec2-GFP. (**C**). Shown are the colocalization of Myo2-GFP with Sec2-mCherry or Sec2GEF-mCherry-CUE at polarized sites (open arrowheads). There is rarely colocalization of Myo2-GFP with Sec2-mCherry or Sec2GEF-mCherry-CUE at non-polarized sites (closed arrowheads). (**D**). Sro7-GFP is well colocalized with Sec2-mCherry and partially colocalized with Sec2GEF-mCherry-CUE at polarized sites (open arrowheads), but there is barely colocalization of Sro7-GFP with either Sec2GEF-mCherry-CUE or Sec2-mCherry at non-polarized sites. Closed arrowheads point to non-colocalized Sro7-GFP with Sec2GEF-mCherry-CUE.

Activation of Sec4 by Sec2GEF-GFP-CUE is also expected to lead to co-localization with downstream Sec4 effectors. The exocyst is normally recruited to secretory vesicles through interactions of Sec4-GTP with the Sec15 subunit (Guo et al., 1999; Salminen and Novick, 1989). We examined cells co-expressing Sec2GEF-GFP-CUE or Sec2-GFP and the exocyst subunit Sec8-mCherry. In both strains, nearly all the GFP structures at polarized sites were also labeled with Sec8-mCherry, however the Sec2GEF-GFP-CUE structures observed at non-polarized sites were labeled to a lower extent with Sec8-mCherry (Fig. 3B). Similar results were observed for cells co-expressing Sec2GEF-GFP-CUE or Sec2-GFP and the exocyst subunit Sec15-mCherry (Supplemental Figure S4A).

Most exocyst subunits are delivered to polarized sites by riding on secretory vesicles along polarized actin cables, while two subunits Sec3 and Exo70 exhibit polarized localization that is partially insensitive to disruption of the actin cytoskeleton (Boyd et al., 2004; Finger et al., 1998; Hutagalung et al., 2009; Liu and Novick, 2014). We examined cells co-expressing Sec2GEF-GFP-CUE or Sec2-GFP and the exocyst subunit Sec3-mCherry. Approximately 40% of Sec2GEF-GFP-CUE puncta were labeled with Sec3-mCherry near sites of polarized growth while Sec2GEF-GFP-CUE structures observed at non-polarized sites were labeled to an even lower extent with Sec3-mCherry. In many cells a Sec2GEF-GFP-CUE puncta was observed directly adjacent to a Sec3-mCherry structure (Supplemental Figure S4C, bottom panel). In control cells expressing Sec2-GFP both proteins were found well colocalized predominantly at polarized sites (Supplemental Figure S4C). Similar results were observed when we co-expressed Sec2GEF-mCherry-CUE or Sec2-mCherry with Exo70-GFP (Supplemental Figure S4D). These findings indicate that in cells expressing Sec2GEF-CUE vesicles are delivered to polarized exocytic sites but are sometimes held at a short distance from sites marked by Sec3 and Exo70.

We examined the co-localization of Sec2GEF-mCherry-CUE or Sec2-mCherry with Myo2-GFP, a type V myosin protein that is recruited to secretory vesicles by Sec4-GTP (Jin et al., 2011). At polarized sites they exhibited co-localization in both strains, but non-polarized Sec2GEF-mCherry-CUE puncta displayed a lower degree of co-localization with Myo2-GFP (Fig. 3C). In the case of Sro7, a SNARE regulator that binds to Sec4-GTP (Grosshans et al., 2006a), even the polarized Sec2GEF-mCherry-CUE puncta displayed a low degree of Sro7-GFP co-localization and non-polarized Sec2GEF-mCherry-CUE puncta were largely unlabeled by Sro7-GFP (Fig. 3D). Together these results indicate that while Sec2GEF-mCherry-CUE can recruit and activate Sec4, the efficiency of recruitment of downstream Sec4 effectors appears to be somewhat reduced relative to full length Sec2, particularly at non-polarized sites and particularly for Sro7. Successful recruitment of Myo2 would be expected to lead to the movement of structures along actin cables towards polarized sites. This explains why the Sec2GEF-mCherry-CUE structures lacking Myo2-GFP are preferentially found at non-polarized sites.

Sec9 is not a direct effector of Sec4. It is a tSNARE that binds to Sro7 and thereby acts downstream of Sec4 (Brennwald et al., 1994; Grosshans et al., 2006a; Lehman et al., 1999). We co-expressed Sec2GEF-mCherry-CUE or Sec2-mCherry and Sec9-GFP. Sec2-mCherry and Sec9-GFP display extensive co-localization at polarized sites. In contrast, Sec2GEF-mCherry-CUE puncta, either at polarized sites or at non-polarized sites, were predominantly negative for Sec9-GFP co-localization (Supplemental Figure S4B). The low extent of co-localization could reflect inefficiency in the activation of SNARE function in response to Sec2GEF-mCherry-CUE, consistent with the analysis of vesicle tethering indicating an extended lifetime of tethered vesicles.

### Sec2GEF-GFP-CUE expression leads to minor dominant negative growth defects and synthetic negative interactions with a subset of secretory and endocytic mutants

We first evaluated the function of the various Sec2 constructs in a *sec2* null background. When expressed from the *SEC2* promoter or from the *ADH1* promoter integrated at the *URA3* locus, both Sec2GEF-GFP-CUE and Sec2GEF-GFP were able to complement the lethality of a *SEC2* deletion (Fig. 4A). Growth was modestly impaired at 37°C in cells expressing Sec2GEF-GFP-CUE from the *SEC2* promoter at the *URA3* locus. A mutation within the CUE domain (M419D) that blocks interaction with ubiquitin moieties restored normal growth. A modest growth defect was observed at 25°C when Sec2GEF-GFP-CUE was expressed from the stronger *Gal1* promoter in either a *sec2Δ* or *SEC2* background (Fig. 4B). This minor, dominant negative effect was blocked by the M419D mutation in the CUE domain. In total, the results indicate surprisingly mild effects on growth of fusing the Sec2GEF to the CUE domain. Nonetheless, the effects observed were at least partially dominant and could be negated by blocking the interaction of the CUE domain with ubiquitin. The *vps4Δ* mutation results in slow growth at 37°C, however expression of Sec2GEF-GFP-CUE in the *vps4Δ* background does not cause any additional growth inhibition (Fig. 4C). The GFP tag appears to confer stability of the fusion proteins since several fold higher levels of Sec2-GFP were detected relative to unfused Sec2 (Elkind et al., 2000), and different tagged versions of Sec2 or Sec2GEF-CUE constructs exhibit similar expression levels (Supplemental Figure S5A). This is indeed the case since a Sec2GEF construct lacking a GFP tag expressed from the *SEC2* promoter integrated at the *URA3* locus was unable to complement the lethality of a *SEC2* deletion (Supplemental Figure S5B). Sec2GEF expressed from the *SEC2* promoter did not exhibit a detectable level of protein while Sec2GEF-GFP expressed from a parallel construct was readily detected (Supplemental Figure S5C).

**Figure 4.**
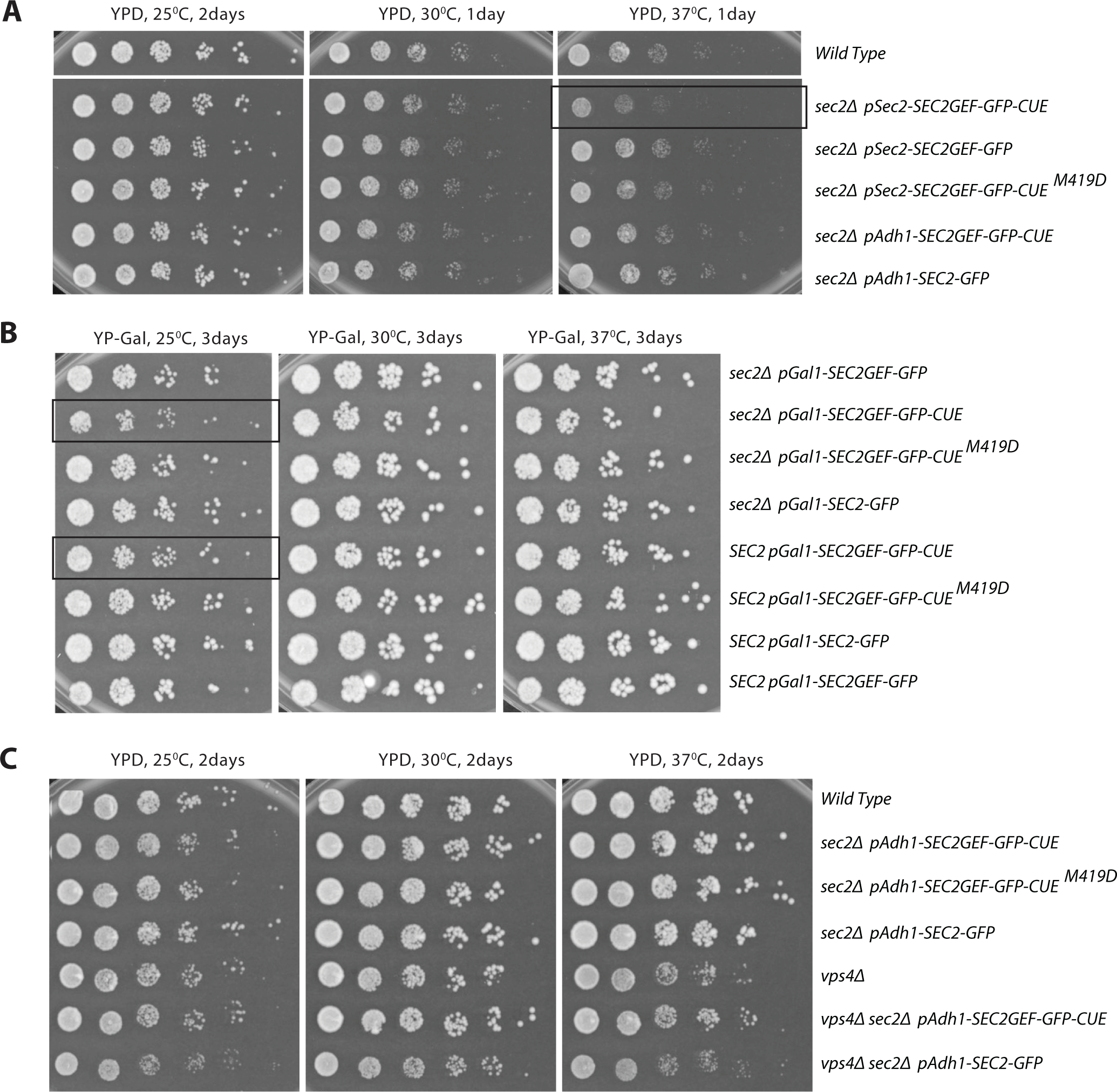
The growth phenotypes of *SEC2GEF-GFP-CUE* mutants expressed from different promoters. (A) Serial dilutions of the overnight cultures from indicated strains were plated on YPD plates and grown for 1 or 2 days at different temperatures as shown. Constructs are expressed from the Adh1 promoter or the Sec2 promoter as indicated. (B) Serial dilutions of the overnight cultures from indicated strains were plated on YP Galactose plates to induce protein expression under the Gal1 promoter and grown for 3 days at different temperatures as shown. Constructs are in either a *sec2Δ* background or are co-expressed with endogenous Sec2 as indicated. Strains exhibiting mild growth defects are framed. (C) Serial dilutions of the overnight cultures from indicated strains were plated on YPD plates and grown for 2 days at different temperatures as shown. Constructs are in a *vps4Δ* or endogenous *VPS4* backgrounds.

Genetic interactions can reveal a partial loss of function and can be informative regarding the site of action of different mutant alleles. We crossed a strain expressing Sec2GEF-GFP-CUE to a variety of different secretory mutants (Table 1). Synthetic negative interactions were observed with a subset of the strains tested, including certain mutations in the exocyst complex (*sec5-24, sec8-9,* and *sec10-2*, but not *sec6-4)*, a tSNARE (*sec9-4*), a SNARE regulator (*sro7Δ*, but not *sec1-1*) and Rab GDI (*sec19-1*). No interaction was seen with the ER mutant *sec12-4*. These results are consistent with an inhibitory effect of Sec2GEF-GFP-CUE on the final stage of the secretory pathway. This effect is not strongly growth limiting on its own but becomes more apparent in a sensitized genetic background.

We also tested the synthetic growth phenotype of Sec2GEF-GFP-CUE with several different endocytic mutants (Table 1). Synthetic negative interactions were observed with a deletion of the endocytic Rab, *ypt51Δ* or its GEF, *vps9Δ*, but no interaction was seen with two endocytic cargo receptor mutants, *syp1Δ or ede1Δ*.

### Sec2GEF-GFP-CUE expression delays the Snc1 cycle at a late stage of the secretory pathway and leads to the accumulation of secretory vesicles

The exocytic vSNARE, Snc1, cycles continuously from secretory vesicles, to the plasma membrane of the bud, then into endocytic vesicles and the Golgi before it is repackaged into a new round of secretory vesicles (Lewis et al., 2000). A slowing of any stage of this cycle leads to a localized build-up of Snc1, even if it is not rate limiting for growth. We co-expressed GFP-Snc1 with Sec2GEF-mCherry-CUE or Sec2-mCherry. In cells expressing Sec2-mCherry, as in wild type cells, GFP-Snc1 was found predominantly at the plasma membrane, preferentially enriched in the bud. In addition, several internal puncta representing endocytic compartments and Golgi were observed in most cells (Fig. 5A). In 45% of cells expressing Sec2GEF-mCherry-CUE a prominent patch of GFP-Snc1 was seen, often in small buds or near the neck of large-budded cells (Fig. 5A). The formation of a patch of GFP-Snc1 near sites of polarized growth is consistent with a delay in the Snc1 cycle at a late stage of the secretory pathway.

**Figure 5.**
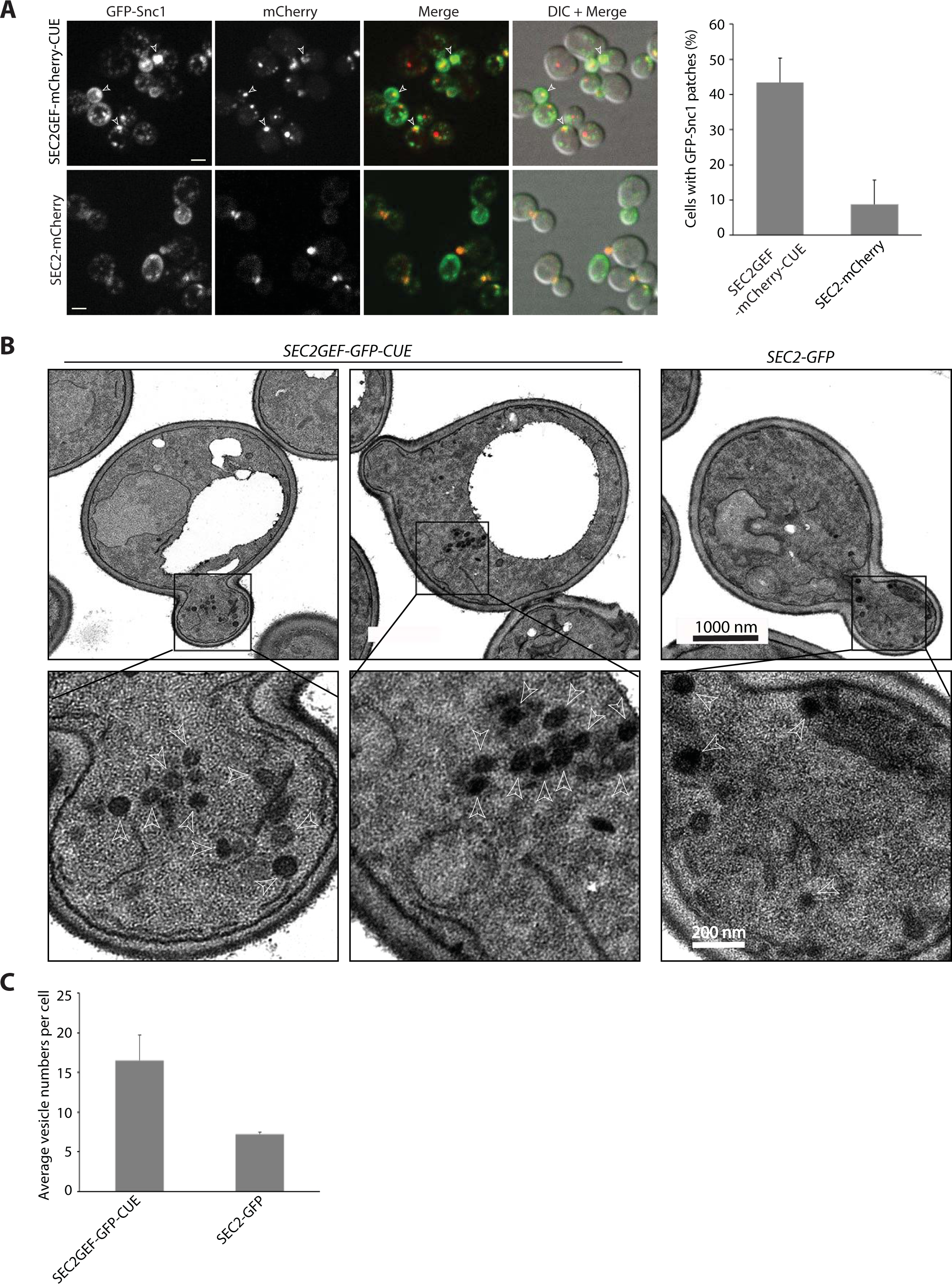
The *SEC2GEF-CUE* mutant has mild secretion defects shown by GFP-Snc1 localization and EM (**A).** Left panel shows representative fluorescence images of GFP-Snc1, the mCherry channel, a merge of the two channels and overlay DIC with merged images in *SEC2-mCherry* or *SEC2GEF-mCherry-CUE* cells grown to early log phase in SC medium at 25°C. Open arrowheads point to accumulated GFP-Snc1 patches. Bars, 5μm. The percentage of cells that contained GFP-Snc1 patches in indicated strains was quantified (right panel). The error bars represent the SD from three independent experiments. (**B)**. Thin-section EM images from the strains indicated were prepared as described in Materials and Methods. The cells were fixed in potassium permanganate. Representative images at 10000x magnification are shown on top panels. The Expanded boxed regions from top panels are shown on bottom panels. Arrowheads indicate vesicles and Scale bars are shown as indicated. (**C)**. Quantitation of secretory vesicles per cell are shown. Over 100 cells were analyzed for each strain.

Thin section electron microscopy analysis revealed the presence of clusters of 80 nm diameter vesicles in cells expressing Sec2GEF-GFP-CUE (Fig. 5B). These clusters were observed in both the bud as well as in the mother cell, consistent with the appearance of the Sec2GEF-GFP-CUE puncta by fluorescence microscopy. Cells expressing Sec2-GFP exhibited isolated 80 nm diameter vesicles which were typically at the bud tip or near the neck. Quantitative analysis demonstrated approximately twice as many vesicles per section in cells expressing Sec2GEF-GFP-CUE relative to cells expressing Sec2-GFP (Fig. 5C).

Despite the accumulation of vesicles, a direct assay of the export of the cell wall glucanase Bgl2 did not reveal a significant intracellular accumulation (Supplemental Figure S6). To explore the possibility of a subtle defect on the secretory pathway we analyzed the effects of Sec2GEF-GFP-CUE expression in a sensitized *sro7Δ* background. The *sro7Δ* single mutant exhibited a minor intracellular accumulation of Bgl2, however the *sro7Δ SEC2GEF-GFP-CUE* double mutant had a much stronger defect (Supplemental Figure S7A). An intermediate result was seen in the *sro7Δ SEC2-GFP* control. Similarly thin section EM showed a much greater accumulation of secretory vesicles in the *sro7Δ SEC2GEF-GFP-CUE* double mutant than in the *sro7Δ* single mutant (Supplemental Figure S7B, S7C). In total, the results support the conclusion that Sec2GEF-GFP-CUE expression leads to a minor defect at the final stage of the secretory pathway that can be exacerbated in a sensitized background.

### Endocytosis of Mup1-GFP is slowed in Sec2GEF-mCherry-CUE expressing cells with a transient association of Sec2GEF-mCherry-CUE with internalized Mup1-GFP

The engineered recruitment of exocytic components onto endocytic membranes raised the possibility of impaired endocytosis. We assessed the rate of endocytosis using the methionine permease, Mup1. In the absence of methionine in the growth medium, this permease resides in the plasma membrane, however within minutes of methionine addition Mup1 is ubiquitinated, internalized into endosomes and delivered to the vacuole for degradation (Guiney et al., 2016). We followed Mup1-GFP through a time course from 3-63 minutes after addition of 20ug/ml methionine. In cells expressing either Sec2GEF-mCherry-CUE or Sec2-mCherry, Mup1-GFP was observed exclusively at the plasma membrane before methionine addition and following methionine addition it was delivered from the plasma membrane to the vacuole (Fig. 6A and Supplemental Figure S8). Rectangular regions of the plasma membrane were selected for quantitation of Mup1-GFP fluorescence intensity. The rate of clearance from the plasma membrane was significantly slowed in Sec2GEF-mCherry-CUE cells relative to Sec2-mCherry cells (Fig. 6B, Supplemental movie S7, S8) as was delivery to the vacuole (Supplemental Figure S8B). Transient co-localization of Mup1-GFP with Sec2GEF-mCherry-CUE was observed. As shown by still frame images in Figure 6C, the two signals either colocalized completely (indicated by open arrowhead) or partially overlapped with each other (indicated by closed arrowhead), suggesting these two proteins might transiently exist within the same compartment rather than directly binding to each other.

**Figure 6.**
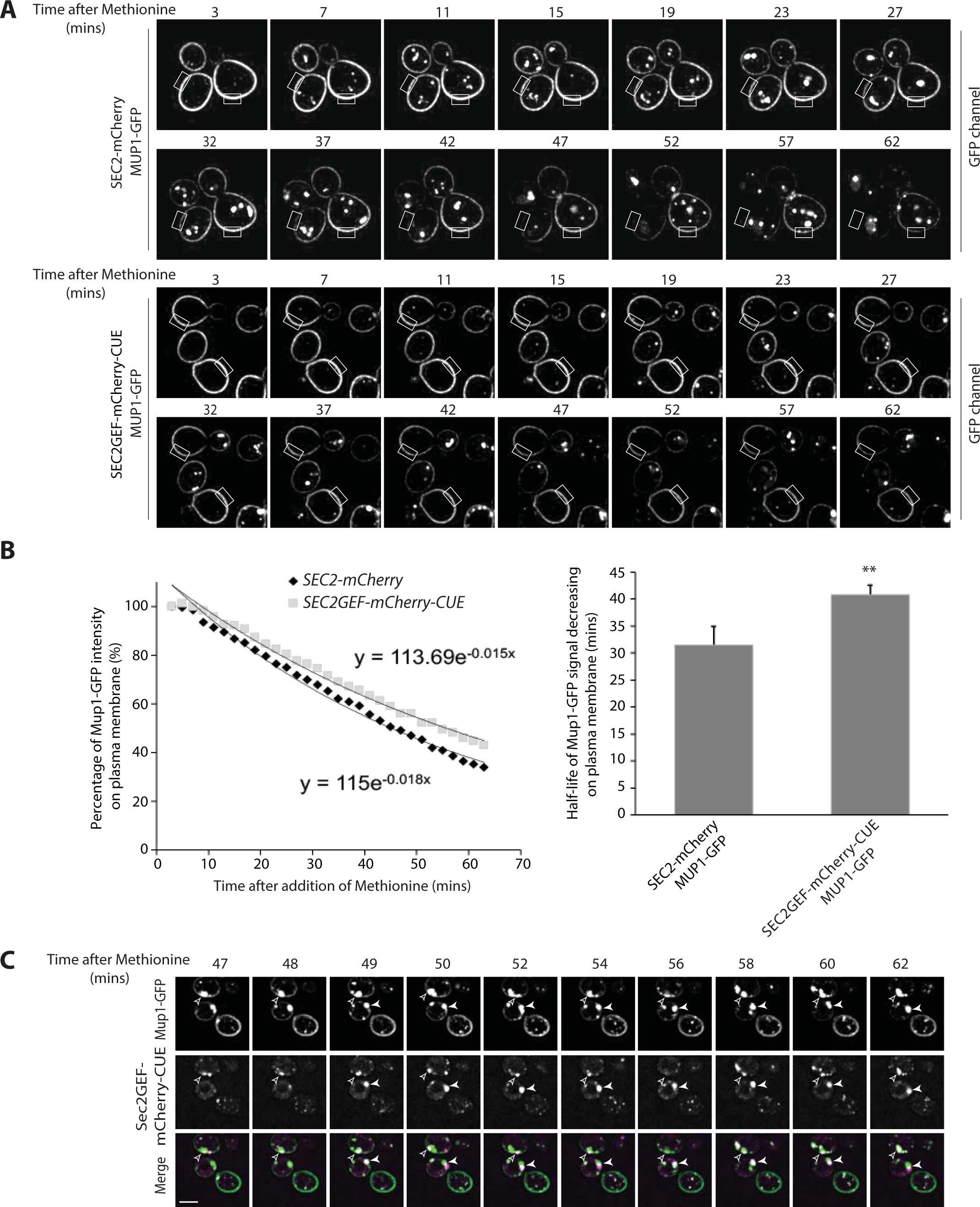
Endocytosis is slow in *SEC2GEF-mCherry-CUE* cells and following internalization Mup1-GFP transiently colocalizes with Sec2GEF-mCherry-CUE. (**A**). Live-cell fluorescence microscopy of Mup1-GFP expressed from the chromosomal locus after the addition of 20μg/ml methionine to stimulate internalization of Mup1-GFP from the plasma membrane and transport to the vacuole lumen in *SEC2-mCherry* (upper panel) or *SEC2GEF-mCherry-CUE* (lower panel) cells (also see Movie S7 and Movie S8). Representative images at indicated time point have been displayed and fluorescence intensity for the rectangle selection area were used for quantitation. (**B**). Relative fluorescence intensity of Mup1-GFP at the plasma membrane was quantified and plotted over time (left panel). Best-fit curves represent a single-exponential decay with one rate of loss from the plasma membrane. The half-times of Mup1-GFP signal decrease on the plasma membrane were 31.5±3.4 s (*SEC2-mCherry*, n=7) and 40.8 ± 1.7 s (*SEC2GEF-mCherry-CUE*, n=7) as shown on right panel. **, *P* < 0.01. (**C**). Still frames of representative cells during Mup1-GFP endocytosis at indicated time points following the addition of 20μg/ml methionine in the *SEC2GEF-mCherry-CUE* strain. Open and closed arrowheads indicate concentrated Mup1-GFP puncta adjacent to the vacuole at which Sec2GEF-mCherry-CUE completely colocalizes or partially overlaps with it. Bar, 5 μm.

## Discussion

We have explored the effects of fusing the catalytic domain of an exocytic Rab GEF onto a CUE endocytic localization domain. Our expectation was that this construct would be recruited to endocytic membranes, which would in turn lead to the ectopic recruitment of Sec4 and its effectors. If Rabs specify the direction of vesicular transport, this could potentially lead to fusion of endocytic compartments with the plasma membrane rather than normal passage along the endocytic pathway. We observed localization of the Sec2GEF-GFP-CUE fusion protein to prominent cytoplasmic puncta that was strongly dependent upon the ubiquitin-binding activity of the CUE domain. Electron microscopy showed that these puncta represent aberrant clusters of 80 nm vesicles. This implies that the fusion protein associates with vesicles containing ubiquitinated endocytic cargo and this was confirmed by the observation of transient colocalization with internalized Mup1. In a *vps4Δ* mutant, in which turnover of ubiquitinated endocytosed proteins is blocked, we observed increased colocalization of Sec2GEF-GFP-CUE with endogenous Vps9, its target Rab, Ypt51, a late endosomal marker Vps8 and a probe for PI3P. As predicted, this triggers recruitment of Sec4 and at least some of its downstream effectors, including the myosin motor, Myo2, resulting in their delivery to polarized sites. Thus, the data suggest that we had in fact directed an exocytic Rab and exocytic machinery onto endocytic membranes, as intended. Nonetheless, the phenotypic effects we observed were surprisingly mild; cell growth and protein secretion were nearly normal. These results imply that Golgi-derived secretory vesicles carrying exocytic cargo were still able to successfully engage the Myo2 myosin motor, the exocyst vesicle tether and the SNARE-dependent fusion machinery, despite the substantial misdirection of Sec4 to endocytic membranes.

A clue to one possible basis for the surprising resilience of the exocytic pathway came from our analysis of the Sec2GEF-GFP construct that we used as a control. This fusion protein, lacking all known Sec2 localization domains was nonetheless able to localize to sites of polarized growth in a small fraction of the cells. Importantly, this residual localization increased upon overexpression of Sec4, suggesting that the Sec2GEF-GFP construct can be recruited to vesicles by Sec4. A downstream effector with an independent localization mechanism, such as the exocyst (Shen et al., 2013), might promote the vesicle association of Sec4 by blocking GDI-mediated extraction, as has been suggested for several other Rabs (Aivazian et al., 2006; Cabrera and Ungermann, 2013). While this mechanism is inefficient, as demonstrated by the low extent of polarized Sec2GEF-GFP localization, it is apparently sufficient to maintain secretory function and normal growth when aided by the increased stability resulting from fusion to GFP. In the case of the Sec2GEF-GFP-CUE construct, this secretory vesicle localization pathway would act in competition with the CUE-domain dependent endocytic localization pathway, to promote secretion and viability. Pertinent to this possibility, we found that deletion of *VPS4* increases recruitment of Sec2GEF-GFP-CUE to enlarged endosomes, but does not impair growth.

Another possible explanation for the mild effects of Sec2GEF-GFP-CUE expression is suggested by the proposal that the TGN serves the role of the early endosome in yeast (Day et al., 2018). By this proposal, endocytic vesicles fuse with the TGN, then vesicles derived from the TGN carrying endocytic cargo are directed to the late endosome. Vps9 is thought to be initially recruited to the TGN, and then incorporated into these vesicles destined for the late endosome (Nagano et al., 2019). If Sec2GEF-GFP-CUE were similarly recruited to the TGN by ubiquitinated endocytic proteins it could potentially help carry secretory cargo proteins destined for the cell surface from the TGN, mimicking the normal exocytic route. This possibility would also explain the observed delay in the delivery of endocytic proteins to the vacuole in Sec2GEF-GFP-CUE cells, as some fraction would be recycled from the TGN back to the cell surface instead of to the late endosome. While we observed no colocalization of Sec2GEF-GFP-CUE with a TGN marker, it is hard to exclude a transient association. It is important to note that our construct contained only the CUE domain of Vps9, while full length Vps9 is thought to also rely on an interaction with Arf1 for TGN recruitment. It is not clear how the lack of an Arf1 binding domain might affect the site of recruitment of Sec2GEF-GFP-CUE.

The association of Sec2GEF-GFP-CUE with endocytic vesicles was not completely benign. We observed weak, dominant negative effects on growth. These effects were relieved by a mutation within the CUE domain that blocks ubiquitin binding, implying that the growth inhibitory mechanism of Sec2GEF-GFP-CUE requires its association with endocytic cargo. Furthermore, delivery of Mup1-GFP to the vacuole was slowed in Sec2GEF-GFP-CUE cells and synthetic negative effects on growth were observed in combination with *vps9Δ* or *ypt51Δ*. The prominent Sec2GEF-GFP-CUE puncta were largely static and did not appear to fuse with the plasma membrane at an appreciable rate, however as these bright puncta represent vesicle clusters, individual vesicles may go on to fuse with the plasma membrane even as the cluster appears static. While most of these puncta acquired Sec4 and its effectors, including the exocyst and Myo2, others failed to acquire Myo2 and were therefore not delivered to polarized sites. Many also failed to acquire the effector, Sro7 and appeared unable to efficiently assemble a SNARE complex with the tSNARE Sec9 despite the presence of the vSNARE Snc1. Although these compartments carried both Sec4 and the Snc1 vSNARE, they were delayed in fusion with the plasma membrane, implying the requirement for an additional component for fusion competence. We speculate that this component could be the phosphoinositide PI(4)P. This lipid is made in the TGN and incorporated into secretory vesicles, but then depleted during vesicle maturation (Ling et al., 2014; Mizuno-Yamasaki et al., 2010; Walch-Solimena and Novick, 1999). While PI(4)P is known to promote Sec2 recruitment (Mizuno-Yamasaki et al., 2010) it might have additional functions related to secretory vesicle function. Endocytic vesicles, in contrast, form PI(3)P which acts to promote the formation of late endosomes (Peterson et al., 1999). We observed significant colocalization of Sec2-GFP-CUE with a PI3P probe in *vps4Δ* cells.

In total, our results indicate that the directionality of vesicular transport cannot be reduced to simply the presence of a particular Rab, but probably involves the interplay between the Rab, its effectors and other vesicular components.

## Materials and methods

### Plasmid and yeast strain construction

The yeast strains and plasmids used in this study are listed in Supplemental Tables SI and SII. To generate the overexpression CEN vector NRB1680 (*pGal1-SEC2GEF-GFP-CUE*), Vps9-CUE (aa408-451) was PCR amplified and by recombination inserted into NRB996 (*pGal-SEC2GEF-GFP, CEN*). Site-directed mutagenesis PCR was used to mutate the M419 codon to Aspartic Acid in the CUE domain to make vector NRB1681 (*pGal1-SEC2GEF-GFP-CUE^M419D^*). To generate the hemiZap vectors NRB1682, NRB1683, NRB1684 and NRB1685, fragments encoding Sec2GEF-GFP, Sec2GEF-GFP-CUE, Sec2GEF-GFP-CUE^M419D^ and full-length Sec2-GFP were PCR amplified from NRB996, NRB1680, NRB1681 and NRB994, then inserted into *Xba*I/*Sal*I double digested pRS306-ADH1t vector. NRB994 and NRB996 were made as previously described in (Elkind et al., 2000). For mCherry or Neongreen tagged versions of Sec2 allele, fragments encoding mCherry or Neongreen were amplified from template NRB1347 (*pJC1-pADH1-mCherry-SEC4*) or NHB0845 (*pFA6a-NeonGreen-His5*, kindly provided by Nan Hao at UCSD) and swapped with the GFP tag in NRB1682, NRB1683, NRB1685 by Gap repair recombination to generate plasmids NRB1686, NRB1687, NRB1688 or NRB1689, NRB1690. Similar approaches were used to generate Sec2 constructs expressed from the *SEC2* promoter. *SEC2* 5′-noncoding sequence upstream of its ORF (770 base pairs) was amplified and switched with the *ADH1* promoter in NRB1682, NRB1683, NRB1684 by Gap repair to generate NRB1691, NRB1692, NRB1693 constructs. The *SEC2* 5′-noncoding sequence upstream of its ORF (770 base pairs) and the first 480 nt coding sequence were amplified and subcloned into *Sac*I/*Xho*I cut plasmid NRB1691 to generate NRB1703. Similarly, The *SEC2* 5′-noncoding sequence upstream of its ORF (770 base pairs) and the first 786nt coding sequence were amplified and subcloned into *Sac*I/*Afl*II cut plasmid NRB1685 to generate NRB1704.

Since *SEC2* is essential, it was necessary to delete the genomic copy of *SEC2* in the diploid strain NY1523. NY3374(*MAT*α*/a, SEC2/sec2Δ::his5*^+^) was generated by replacing one copy of the *SEC2* coding sequence with the *Schizosaccharomyces pombe his5*^+^ gene module using the PCR-mediated gene deletion method (Longtine et al., 1998). To construct strains expressing various *sec2* alleles as the sole copy in yeast, HemiZAP *sec2-*expressing plasmids were digested with a unique enzyme (see Supplemental Table SII) and introduced into yeast diploid cells (NY3374) at the *URA3* locus by homologous recombination. After sporulation and tetrad dissection, the spores were analyzed and we selected representatives that carried only the mutant *sec2 allele* with a C-terminal tag. For *GAL1* promoter driven overexpression constructs of *SEC2*, CEN plasmids NRB996, NRB1680, NRB1681 and NRB994 were directly transformed into wild type yeast cells NY1210 and grown in SC-Ura3 with 2% Galactose condition to induce *sec2* allele overexpression in the presence of wild type untagged *SEC2*. Alternatively these CEN plasmids were transformed into a diploid yeast strain (NY3374) and after sporulation and tetrad dissection the haploids with only one copy of the mutant *sec2* allele on a plasmid were selected.

Two *SEC4* integration plasmids were used. One is tagged with mCherry at its N-terminus and overexpressed from the strong *ADH1* promoter (NRB1347). Another one is tagged with GFP at its N-terminus and expressed under its own promoter (NRB1694). It was subcloned from NRB1312 (*pRS306-pSec4-GFP-SEC4-tSEC4*) into NRB1695 (*pRS305-myo2-GFP*) using *Xho*I*/Sac*I. NRB1347 was linearized with *Afl*II to integrate into *LEU2* locus while NRB1694 was linearized with *Sph*I to integrate into the *SEC4* locus. To generate the hemizap *pRS305-myo2-GFP* vector (NRB1695), a C-terminal fragment of *MYO2* (3189-4722bp) was amplified by PCR from gDNA and subcloned into the *pRS305-GFP* plasmid with *Xho*I and *Sac*II. The plasmid was linearized using *Stu*I and integrated into the *MYO2* locus. Similar approaches were used to generate *EXO70*, *SEC3*, *SEC8*, *SEC15* and *VPS9* HemiZAP constructs (NRB1565, NRB1329, NRB1460, NRB1697, NRB1696).

To generate the hemizap *GFP-SEC9-LEU2* vector (NRB1698), a fragment encoding the *SEC9* promoter 1026bp, the GFP module 738bp and the *SEC9* N-terminal 705bp ORF were amplified by PCR from gDNA or a common plasmid and sequentially subcloned into plasmid NRB1631 using *Xho*I/X*ma*I, *Xma*I/*Xba*I and *Xba*I/*Pst*I to replace the sequence of *pSec5-Tos2(2-74)-Sec5N*. The final plasmid was linearized with *Bgl*II and transformed into yeast at the *SEC9* locus. To generate the mCherry-Ypt51 integration vector (NRB1700), the *YPT51* ORF sequence was amplified by PCR from gDNA and subcloned into NRB1347 to replace the *SEC4* sequence using *BamH*I/*Sal*I. This plasmid was linearized using *Afl*II and integrated into the *LEU2* locus. To switch the auxotrophic marker of the Vps8-mCherry integration vector (NRB1699), the *VPS8-6xmCherry* sequence was excised from *YIplac211-VPS8-6xmCherry* (generous gift from B. Glick) (Losev et al., 2006) and subcloned into pJC1 vector by *Hind*III/*Sac*I. This plasmid was linearized using *Bcl*I and integrated into the *VPS8* locus. Similarly the *SEC7-6xdsRed* sequence was excised from NRB1338 (*YIplac204-SEC7-6xdsRed)* (Losev et al., 2006) and subcloned into pJC1 vector by *BstAP*I/*Pvu*I. This plasmid (NRB1702) was linearized using *Age*I and integrated into the *LEU2* locus. To switch the auxotrophic marker of the mRFP-FYVE(EEA1) integration vector (NRB1706), the *mRFP-FYVE(EEA1)* sequence was excised from *pRS416-mRFP-FYVE(EEA1)* (generous gift from Seth Field) and subcloned into pRS305 vector by *Xho*II/*Sac*I. This plasmid was linearized using *Afl*II and integrated into the *LEU2* locus. Similarly the *FAPP1-PH-Cyc1_term_* sequence was subcloned from *pRS306-mCherry-FAPP1-PH* (NRB1444) (Mizuno-Yamasaki et al., 2010) into *YIplac128-pADH1-mCherry* vector by *BamH*I/*Eag*I. This plasmid (NRB1705) was linearized using *Afl*II and integrated into the *LEU2* locus. To generate the HemiZap vector *pRS305-mup1-GFP* (NRB1701), first a fragment containing GFP and the *Cyc1* terminator was excised from NRB1685 with *Nhe*I and *Not*I and used to replace the mCherry-Cyc1_ter_ part in NRB1699 (*pRS305-vps8-mcherryx6*), then a C-terminal fragment of *Mup1*(740-1723bp) was amplified by PCR and subcloned into above plasmid with *Sal*I and *Nhe*I. This plasmid was linearized with *Bsu36*I and transformed into yeast at the *MUP1* locus.

All the other yeast strains were constructed by recombination to delete the gene or tag the gene chromosomally (Longtine et al., 1998) or by a genetic cross to select a haploid of the appropriate genotype.

### Growth test

Yeast cells were grown overnight in yeast peptone dextrose (YPD) or synthetic complete (SC) dropout medium to stationary phase. Cells were washed once with sterile water and spotted on YPD or SC dropout plates in fivefold serial dilutions starting with an OD_600_ of 0.2. For GAL induction experiments, yeast cells were grown in YP medium containing 2% raffinose overnight and then spotted on YPGAL plates. Plates were incubated at the specified temperature for the indicated time.

### Cell lysate extracts and Immunoblotting

Yeast cells were grown to OD_600_ ∼ 1 at 25°C. For each strain an equal amount of cells were harvested and lysed by a rapid alkaline lysis procedure as previously described (Westfall et al., 2008). Proteins were subjected to Western blot analysis with mouse monoclonal anti-GFP antibody (1:2,000 dilution, Roche), rabbit polyclonal anti-Sec2 antibody (1:2000 dilution, lab made) or mouse monoclonal anti-Pgk1(phosphoglycerate kinase) antiboty as loading control (1:5000 dilution, Invitrogen).

### Bgl2 secretion assay

The Bgl2 secretion assay was performed as described by (Kozminski et al., 2006) with minor modification. In brief, yeast cells (20 ml) were grown at 25°C in YPD overnight to early log phase (0.4-0.8 OD_600_/ml). About 5 OD_600_ units of cells were harvested by centrifugation at 900 × *g* for 5 min. Three different sets of cultures were prepared for each strain. Cell pellets were resuspended in 1 ml of ice-cold 10 mM NaN_3_ and 10 mM NaF, followed by a 10-min incubation on ice. The suspension was transferred to microfuge tubes, pelleted, and resuspended in 1 ml of fresh prespheroplasting buffer (100 mM Tris-HCl, pH 9.4, 50 mM β-mercaptoethanol, 10 mM NaN_3_, and 10 mM NaF). After a 15-min incubation on ice, cells were pelleted and washed with 0.5 ml of spheroplast buffer (50 mM KH_2_PO_4_-KOH, pH 7.0, 1.4 M sorbitol, and 10 mM NaN_3_).

Cells were resuspended in 1 ml of spheroplast buffer containing 167 μg/ml zymolyase 100T (Nacasai Tesque Inc.). Cells were incubated in a 37°C water bath for 30 min. Spheroplasts were then pelleted at 5000 × *g* for 10 min, and 100 μl of the supernatant was transferred into a new tube and mixed with 50 μl of 3× SDS sample buffer. This represents the external pool. All the remaining supernatant was removed, and the spheroplast pellet was rinsed once with 1 ml of spheroplast buffer and then resuspended in 100 μl of 2× SDS sample buffer. This represents the internal pool. Proper amount of each internal pool sample and external pool sample were loaded onto a 12% SDS–PAGE gel. Bgl2 was visualized by Western blotting with anti-Bgl2 rabbit polyclonal antibody at 1:5000 dilution (provided by R. Schekman, University of California, Berkeley, CA). The amount of Bgl2 in both internal and external pools was determined by ImageJ software. The fraction of internal Bgl2 accumulated was calculated by: Int/(Int+Ext).

### Electron Microscopy

Yeast cells expressing full length Sec2-GFP or Sec2GEF-GFP-CUE from the *ADH1* promoter in the *sro7Δ::LEU2* background (NY3398 and NY3395) as well as control cell *sro7Δ::LEU2* (NY2599) were grown at 25 °C in YPD to an OD_600 nm_ of ∼0.5 and then processed for electron microscopy as previously described (Chen et al., 2012). Alternatively, yeast cells expressing full length Sec2-GFP or Sec2GEF-GFP-CUE from the *ADH1* promoter in the *wild-type* background (NY3384 and NY3381) were grown at 25 °C in YPD to an OD_600 nm_ of ∼0.5 and electron microscopy was performed as described (Cui et al., 2019). Briefly, cells were fixed in potassium permanganate and embedded in Spurr’s resin. After polymerization, 55- to 60-nm thin sections were cut using an Ultracut ultramicrotome (Leica Microsystems), transferred onto formvar carbon-coated copper grids, and stained before viewing. For both preparation conditions, the images were acquired using a transmission electron microscope (Tecnai G2 Spirit; FEI) equipped with a CCD camera (UltraScan 4000; Gatan).

### Fluorescence microscopy and quantitative localization analysis

Yeast strains harboring GFP, mCherry or dsRed tags were grown overnight to early log phase (OD_600_ 0.4–0.6) in selective SC medium at 25°C. Live cells (500 μl) were pelleted by centrifugation at 5000 rpm for 1 min and were resuspended in 50 μl of medium. 5 μ of the cell suspension was spotted on a glass slide with coverslip. The cells were examined on two different microscopes, typically on an Axiophot 2 upright microscope (Carl Zeiss microimaging, Inc) with 100x Plan apochromatic oil-immersion objective lens with NA 1.3 and 100W xenon excitation lamp and with a cool-CCD camera (model ORCA ER; Hamamatsu). As specified, some fluorescence images were acquired on a Yokagawa spinning-disk confocal microscopy system (Zeiss Carl Observer Z1) with a 100× oil immersion objective lens (Plan Apochromat 100×/1.4 NA oil DIC lens; Carl Zeiss) equipped with an electron multiplying CCD camera (QuantEM 512SC; Photometrics). Excitation of GFP or mCherry/dsRed was achieved using 488-nm argon and 568-nm argon/krypton lasers, respectively. For each sample, a z-stack with a 200-nm slice distance was generated. Images were analyzed using velocity or AxioVision software 4.8 (Carl Zeiss).

To quantify the localization of Sec2 or Sec2GEF-CUE, about 50 cells were examined for each condition and at least three independent clones were tested for each strain to calculate the SD. When the GFP/mCherry signal was mainly detected at the tips of small/medium budded cells, and at the bud necks of large-budded cells, these cells were scored as polarized, otherwise, when the GFP/mCherry signal was mainly detected as puncta inside the cytosol, away from the bud tip or bud neck area, these cells were scored as non-polarized. Those that containing mostly diffuse GFP or mCherry signal were not counted.

### Time-lapse imaging and analysis of Sec2 or Sec2GEF-CUE containing vesicles

The time-lapse fluorescence imaging was conducted on yeast strains expressing full length Sec2-Neongreen or Sec2GEF-Neongreen-CUE. Cells were grown overnight in SC medium to early log phase (0.3-0.5 OD600), and then were applied to a 35-mm MatTek dish (Ashland) pretreated with 0.05 mg/ml Concanavalin A (EY Laboratories, Inc.) for imaging on a Yokogawa W1-SoRa spinning disc confocal mounted on a Nikon Ti2-E microscope body. Images were acquired with a 100x Plan Apo Lambda 1.35 NA silicone immersion objective. A Nikon LUNF-XL 6-line (405, 445, 488, 520, 561, and 640 nm) laser engine, Prime 95B back-thinned sCMOS camera (Teledyne Photometrics), piezo Z-stage (Mad City Labs), were used to acquire images via NIS Elements software. Images were taken with the SoRa mode with 2.8x additional magnification, 100ms exposure time, and 20%-40% 488-nm laser power. To train an AI enhancement model with the enhance.ai module in NIS Elements software a set of over 100 image pairs of these cells were acquired with the first image acquired with the above imaging settings with the second was acquired with 80% 488-nm laser power and 150ms exposure time. Training of the AI model proceeded to a training loss of .136, which was largely due to the fact that there was movement between the two image frames as the sample was unfixed. The enhance.ai model was incorporated into a GA3 workflow, which allowed us to batch apply it to all images.

Individual vesicles that have the expected intensity and display a clear moving track were identified by eye. The life-time for vesicle puncta was quantified by kymograph in ImageJ v2.1. Briefly, a timelapse image was max-projected in time to create a new image in which the movement of individual vesicles could be detected as lines. These lines were traced and added to the Region Of Interest (ROI) manager. In the ROI Manager, selected segmented lines were placed in the original image stacks, then the Reslice Stack command was used to obtain kymograph images of the objects where the y-axis distance represents the life-time of the vesicles. Due to the time duration of the imaging, some of the vesicles tracked remained visible until the end of imaging, therefore the lifetimes were grouped into three ranges and the vesicle dynamics can be compared by their distribution within the different ranges.

### FRAP experiment and image analysis

The live-cell preparation and time-lapse imaging after photobleaching for FRAP experiments were conducted with the same spinning disc confocal microscope system as described above. Photobleaching of the entire bud was performed using an OptiMicroScan X-Y Galvo system with a 1-ms dwell time and the laser intensity was set to 10% from a directly coupled 20mW 405nm laser. The bleach duration was chosen by determining the minimal time needed to completely bleach the fluorescent signal. To increase imaging speed and avoid photobleaching, images were taken using a 256 × 256 pixel ROI. Three prebleached images were acquired without a delay, and postbleaching images were taken continuously for 30s and then with 1s intervals for a subsequent 1min. In the vesicle tracking experiments after photobleaching, the movies were taken in a z stack of 5 planes (covering 2.5um), each with 100ms exposure time from 20-40% 488-nm laser power.

ROIs for the bleached bud and the whole cell as well as a rectangular background ROI in a dark extracellular region were drawn by hand and moved into the ROI Manager. The Time Measurement tool within NIS Elements was used to quantify the intensity of the bleached ROIs as well as the entire cell after the background intensity was subtracted from the images. The fluorescence recovery within the bleached bud relative to the total signal within the whole cell was plotted over time, and the prebleached signal intensity was normalized as 100%. The pixel intensity change of a nonbleached cell over time was collected to adjust for the photobleaching effects of imaging.

Sec2-Neongreen or Sec2GEF-Neongreen-CUE positive vesicles entering the bud after FRAP were tracked by eye until one was observed tethering to the plasma membrane or relatively stabilized around the plasma membrane area. Only those vesicles free of adjacent vesicles or overlapping vesicles throughout the tracking time were analyzed. If the vesicles disappeared from plasma membrane sites, we checked in other z-stack planes to confirm that they vanished due to fusion and not due to drift out of the field of view in the z direction. Static images were converted to 16-bit and displayed in inverted monochrome maximum projection, then range-adjusted to the minimum and maximum contrast with ImageJ. Due to dramatic dynamic difference of vesicles in wild-type Sec2-Neongreen and Sec2GEF-Neongreen-CUE cells and the relatively low intensity of vesicles, kymograph by line around bud cortex in NIS software showed large variations. Quantitation of the time from appearance to disappearance for individual vesicle puncta was obtained manually for more than 20 cells for each strain.

### Endocytosis of Mup1-GFP and image quantitation

To track endocytosis, we integrated the GFP sequence at the C-terminus of Mup1 in *wild-type, SEC2-mCherry,* or *SEC2GEF-mCherry-CUE* strains. Cells were grown overnight to early log phase (0.3-0.5 OD_600_) in SC medium without Methionine and were applied to a 35-mm MatTek dish pretreated with 0.05 mg/ml Concanavalin A. Cells were quickly rinsed with SC medium without Methionine and then fresh SC medium containing Methionine was added to stimulate endocytosis before conducting time-lapse imaging on a Yokogawa X1 spinning disk confocal system attached to a Ti2-E. The Nikon microscope employed was equipped with a 100x Apo TIRF 1.49 NA objective, Prime 95B back-thinned sCMOS camera and NIS Elements software. Acquisition settings for Mup1-GFP endocytosis were 60min duration, 1-min interval time, 200- ms exposure time, 20% 488nm laser power, one z-axis plane with PFS setting to prevent drifting. The temperature of the immersion oil on the microscopy slide near the sample was ∼24°C.

Image processing and quantitation were performed using the provided NIS Elements software. Rectangular ROIs in the cell plasma membrane were drawn manually to exclude any obvious puncta from inside or next to the ROIs. The Time Measurement tool within NIS Elements was used to quantify the intensity of the plasma membrane in all selected ROIs, and then results were exported to Excel. The intensity at the starting time point was set as 100%, then the percentage of Mup1-GFP intensity residing on the plasma membrane was plotted over time. At least three independent experiments were performed and a minimum number of 50 yeast cells were quantified from each experiment. The averaged intensity decrease for one representative experiment was shown. Best-fit exponential trendlines with 2^nd^ order decay were applied and a τ value, half-life of Mup1-GFP on plasma membrane, was determined. The averaged τ values for Mup1-GFP in *SEC2-mCherry* or *SEC2GEF-mCherry-CUE* were from at least three endocytosis experiments. Probability values (*P* value) were calculated using the Student’s *t* test, and all comparisons with a *P* value < 0.005 were considered statistically significant.

The quantitation of cell percentage containing GFP signal in the vacuole was done manually at two time point 63min and 93min after the addition of methionine. The time-lapse imaging were acquired on a Yokagawa spinning-disk confocal microscopy system (Zeiss Carl Observer Z1) equipped with a 100× oil immersion objective len, an electron multiplying CCD camera (QuantEM 512SC; Photometrics) and AxioVision software 4.8 (Carl Zeiss). Acquisition settings for Mup1-GFP endocytosis were 90min duration, 15-min interval time, 200-ms exposure time, 80% 488nm laser power, and a z stack of 5 planes with a step 0.25um.Three independent experiments were performed and a minimum number of 30 yeast cells were quantified from each experiment.

## Acknowledgements

This work was supported by grants GM35370 and GM08261 to P.N. from the N.I.H. and by the George E. Palade Endowed Chair and Margret Shaw Roberts Fund. We thank Dr. Randy Schekman, University of California at Berkeley, Dr. Nan Hao, University of California at San Diego, Dr. Seth Field, Case Western Reserve University and Dr. Benjamin Glick, University of Chicago for reagents. We also thank Drs. Hua Yuan and David Shen for their assistance in strain construction.

---

**Figure S1.** Vps9-mCherry partially colocalizes with Sec2GEF-GFP-CUE but not with Sec2-GFP. Representative images of Sec2-GFP or Sec2GEF-GFP-CUE, Vps9-mCherry, merged and overlay DIC with merged images are shown on left panel. Closed arrowheads indicate GFP puncta with which Vps9-mCherry is not colocalized. Open arrowheads and arrows point to GFP puncta colocalized with Vps9-mCherry at polarized sites and non-polarized sites, respectively. The localization of Sec2GEF-GFP-CUE or Sec2-GFP and their colocalization with Vps9-mCherry were quantified and shown on the right panel. The error bars represent SD from three independent experiments. Bar, 5 μm.

**Figure S2.** Localization of Sec2GEF-GFP-CUE (Sec2GEF-mCherry-CUE) or Sec2-GFP (Sec2-mCherry) and their colocalization with Sec4, Sec8 or Sec7 in *vps4*Δ strains. **(A).** Confocal fluorescence micrographs of Sec2-GFP or Sec2GEF-GFP-CUE in *vps4*Δ strains. Left panel shows representative GFP fluorescence, DIC and merged images of Sec2-GFP or Sec2GEF-GFP-CUE in *vps4*Δ background, and the quantitation results of GFP puncta localized at polarized sites (open arrowheads) or non-polarized sites (arrows) are shown on right panel. **(B).** Live-cell fluorescence microscopy of *SEC2-mCherry* and *SEC2GEF-mCherry-CUE* strains expressing GFP-Sec4 in *vps4*Δ strains. Left panel shows representative GFP fluorescence, mCherry fluorescence, merged, and DIC overlaid with merged images. The quantification of Sec2-mCherry and Sec2GEF-mCherry-CUE localization at polarized or non-polarized sites and their colocalization with GFP-Sec4 protein are shown on the right panel. Open arrowheads and arrows indicate respectively polarized sites and non-polarized sites at which mCherry colocalizes with GFP-Sec4 protein. **(C-D).** Upper panels show representative images of GFP channel, Sec8-mCherry (C) or Sec7-dsRed (D), merged channels and overlay DIC with merged images in *SEC2-GFP* or *SEC2GEF-GFP-CUE* strains. Open arrowheads indicate polarized sites at which GFP colocalizes with dsRed or mCherry-tagged proteins, while closed arrowheads point to GFP puncta which are not colocalized with the proteins examined. The quantification of Sec2-GFP and Sec2GEF-GFP-CUE colocalization with Sec8-mCherry or Sec7-dsRed are shown on lower panels. In all quantitation graphs the error bars represent the SD from three independent experiments. Scale bar, 5 μm.

**Figure S3.** Colocalization of Sec2GEF-GFP-CUE or Sec2-GFP with mCherry-FAPP1-PH or mRFP-FYVE(EEA1) in *wild type VPS4* (A and C) or *vps4*Δ strains (B and D). (**A) and (B).** Shown on upper panels are representative Sec2-GFP or Sec2GEF-GFP-CUE, mCherry-FAPP1-PH, merged and overlay DIC with merged images. Closed arrowheads point to Sec2-GFP or Sec2GEF-GFP-CUE not colocalizated with mCherry-FAPP1-PH. There is very low percentage of colocalization of mCherry-FAPP1-PH with Sec2GEF-mCherry-CUE in *vps4*Δ strains. The quantification results are shown on the lower panels. **(C) and (D).** Shown on top panels are representative images of the mRFP channel, GFP channel, merged channels and overlay DIC with merged channels. Open arrowheads or closed arrowheads point to respectively colocalized or non-colocalized Sec2-GFP or Sec2GEF-GFP-CUE with mRFP-FYVE(EEA1). There is a significant amount of colocalization of mRFP-FYVE(EEA1) with Sec2GEF-GFP-CUE in *vps4*Δ strains at both polarized and non-polarized sites, but no colocalization of mRFP-FYVE(EEA1) with Sec2-GFP in either strain. The quantification results are shown on bottom panels. The error bars represent SD from three independent experiments. Scale bar, 5 μm.

**Figure S4.** Colocalization of Sec2GEF-GFP-CUE, Sec2GEF-mCherry-CUE, Sec2-GFP or Sec2-mCherry with Sec15-mCherry, GFP-Sec9, Sec3-mCherry or Exo70-GFP. (**A).** Shown on upper panel are representative Sec2-GFP or Sec2GEF-GFP-CUE, Sec15-mCherry, merged and overlay DIC with merged images. Open arrowheads point to Sec15-mCherry colocalization with Sec2-GFP or Sec2GEF-GFP-CUE at polarized sites. The quantification results are shown on the lower panel. (**B**). GFP-Sec9 expressed from the chromosomal locus exhibits little co-localization with Sec2GEF-mCherry-CUE. Upper panel shows representative mCherry-labeled Sec2 or Sec2GEF-CUE, GFP-Sec9, merged and overlay DIC with merged images. Open arrowheads point to its colocalization with Sec2-mCherry or Sec2GEF-mCherry-CUE at polarized sites, while closed arrowheads point to non-colocalized GFP-Sec9 with Sec2GEF-mCherry-CUE. There is a low percentage of colocalization of GFP-Sec9 with Sec2GEF-mCherry-CUE at polarized sites and barely any colocalization with Sec2-mCherry or Sec2GEF-mCherry-CUE at non-polarized sites. The quantification results are shown on lower panels. **(C) and (D).** Shown on top panels are representative images of the mCherry channel, GFP channel, merged channels and overlay DIC with merged channels. Open arrowheads or closed arrowheads point to respectively colocalized or non-colocalized exocyst Sec3 or Exo70 with Sec2 or Sec2GEF-CUE at polarized sites. There is little colocalization of exocyst Sec3 or Exo70 with Sec2 or Sec2GEF-CUE at non-polarized sites. The quantification results are shown on the middle panels. Among the Sec3 or Exo70 structures that fail to colocalize with Sec2GEF-CUE at polarized sites, most are directly adjacent to a Sec2GEF-CUE structure. The percentage of cells containing Sec2GEF-CUE or Sec2 at polarized sites among which Sec3 or Exo70 is either colocalized or adjacent were quantified and shown on bottom panels. The error bars represent SD from three independent experiments. Scale bar, 5 μm.

**Figure S5.** Sec2GEF expressed from the Sec2 promoter is not sufficient to support cell growth while GFP tag and other tags confers stability of the fusion proteins. (**A).** Protein expression level from the GFP-, mCherry- and NeonGreen-tagged Sec2 or Sec2GEF-CUE constructs under ADH1 promoter have been examined by anti-Sec2 western blot. The yeast lysates were extracted from strains NY3381, NY3384, NY3388, NY3390, NY3391 and NY3392, separately. Pgk1 was visualized as a loading control. (**B).** Tetrad dissection analysis for diploid cells with one chromosomal *SEC2* copy replaced with the *HIS3* module and Sec2GEF expressed from the Sec2 promoter integrated at the *ura3* locus. After dissection, replica plating to infer genotype showed that the inviable spores were *HIS3* and were either *URA3* or *ura3*. (**C**). Yeast lysates were extracted separately from strains *wild type* (NY1210), *sec2Δ ura3-52::pSec2-SEC2GEF-GFP-CUE* (NY3385), *sec2Δ ura3-52::pSec2-SEC2GEF-GFP-CUE^M419D^* (NY3386), *sec2Δ ura3-52::pSec2-SEC2GEF-GFP* (NY3387) or *SEC2 ura3-52::pSec2-SEC2GEF* (NY3445). Sec2 protein level was measured using anti-Sec2 western blot.

**Figure S6.** *SEC2GEF-CUE* cells do not exhibit significant secretion defects by Bgl2 secretion assay. (**A)**. Strains *WT*, *SEC2GEF-GFP-CUE, SEC2GEF-GFP-CUE^M419D^, SEC2GEF-GFP, SEC2-GFP* or secretion mutant strain *sec6-4* were grown at 25^0^C in YPD overnight to early-log phase and shifted to 37^0^C for 1hr before harvesting. Internal and external fractions of Bgl2 glucanase were prepared as described in Materials and Methods. Eight-fold more of the internal fraction was loaded relative to the external fraction. Bgl2 was visualized by western blotting using anti-Bgl2 rabbit polyclonal antibody. (**B)**. Normalized quantitation of internal Bgl2 percentage from the blots on panel **A** is shown. Error bars represent SD from three independent experiments.

**Figure S7.** The *SEC2GEF-CUE* mutant has very mild secretion defects alone, yet the defects are more severe in *sro7*Δ cells. (**A).** Strains WT, *sro7Δ, sro7Δ SEC2GEF-GFP-CUE, sro7Δ SEC2-GFP* were grown at 25^0^C in YPD overnight to mid-log phase. Internal and external fractions of Bgl2 glucanase were isolated and analyzed as described in Materials and Methods. Ten-fold more of the internal fraction was loaded relative to the external fraction. Bgl2 was visualized by western blotting using anti-Bgl2 rabbit polyclonal antibody. Normalized quantitation of internal Bgl2 percentage from the blots on the left panel is shown on the right panel. Error bars represent SD, n=3. ns, not significant, * P< 0.05, ** P< 0.01. (**B-C)**. Thin-section EM images from the strains indicated are prepared as described in Materials and Methods. The cells were fixed in cacodylate/glutaraldyhyde and stained with lead citrate/uranyl acetate. Representative images at 10000x magnification are shown. Arrowheads indicate vesicles and Scale bars are shown as indicated. Quantitation of secretory vesicles per cell are shown in panel C. Over 100 cells were analyzed for each strain.

**Figure S8.** Endocytosis followed by Mup1-GFP delivery to the vacuole for degradation is slow in *SEC2GEF-mCherry-CUE* cells. (**A**). Live-cell fluorescence microscopy of Mup1-GFP in *Sec2GEF-mCherry-CUE* (upper panel) and *Sec2-mCherry* cells (lower panel) after the addition of 20μg/ml methionine to stimulate internalization of Mup1-GFP from the plasma membrane and transport to the vacuole lumen. Representative images at indicated time point have been displayed and GFP fluorescence patches in the vacuole were indicated by arrowheads. Scale bar, 2 μm. (**B**). The percentage of cells containing GFP fluorescence in the vacuole at two time points 63min and 93min after the addition of methionine was quantified and plotted. Error bars represent SD from three independent experiments.

**Movies S1 and S2.** Vesicle dynamics as seen in Sec2GEF-Neongreen-CUE labeled cells (S1) and in wild-type Sec2-Neongreen labeled cells (S2). The acquisition was taken using SORA super resolution setting with 40% 488nm laser power and images are processed using AI enhancement. Acquisition time duration is 30sec without interval delay; movie speed, 10 frames per second = 1.7s/s. One z-axis plane fluorescence image was acquired.

**Movies S3 and S4.** Examples of Fluorescence Recovery After Photobleach (FRAP) in entire small buds of *SEC2GEF-Neongreen-CUE* cells (S3) and wild-type *SEC2-Neongreen* cells (S4). Acquisition was taken using regular setting with 40% 488nm laser power. Photobleaching of the entire bud was performed using 10% laser power at 405nm with a 1-ms dwell time and a raster block size of 10 (duration 100ms/point). Acquisition time duration is 30sec without interval delay; movie speed, 10 frames per second = 2s/s. One z-axis plane fluorescence image was acquired.

**Movies S5 and S6.** Show examples of bleached cells used in vesicle tracking studies for wild-type *SEC2-Neongreen* (S5) and *SEC2GEF-Neongreen-CUE* (S6) strains. Acquisition was taken using SORA super resolution setting with 20% 488nm laser power and images were processed using AI enhancement. Photobleaching of the entire bud was performed using 10% laser power at 405nm with a 1-ms dwell time and a raster block size of 10 (duration 100ms/point).

Acquisition time duration is 1min with 1sec interval for *SEC2GEF-Neongreen-CUE* strain and 30sec with 1sec interval for *SEC2-Neongreen* strain since overall signal in *SEC2-Neongreen* strain was weaker; movie speed, 10 frames per second = 10s/s. Five z-axis plane fluorescence images were acquired and merged.

**Movies S7 and S8.** Endocytosis of Mup1-GFP in *SEC2-mCherry* cells (S7) and in *SEC2GEF-mCherry-CUE* cells (S8). The acquisition was started 3min after 20μg/ml methionine addition. Acquisition time duration was 60min with 1min interval; movie speed, 2 frames per second = 2 min/s. One z-axis plane fluorescence image was acquired.

**Table I:** Synthetic growth defects of secretory mutant or endocytic mutant strains when combined with *SEC2GEF-GFP-CUE*.

**Table SI:** Yeast strains.

**Table SII:** Bacterial plasmids.

